# BAZ1A promotes expression of *DUX4-fl* and its lncRNA activator *DBE-T* in facioscapulohumeral muscular dystrophy

**DOI:** 10.64898/2026.06.15.732407

**Authors:** Ning Chang, Takako I. Jones, Peter L. Jones, Charis L. Himeda

## Abstract

Facioscapulohumeral muscular dystrophy (FSHD) is caused by incomplete epigenetic silencing of a D4Z4 macrosatellite array, leading to pathogenic misexpression of *DUX4* in skeletal muscle. Therapeutic development of small molecule drugs for FSHD has been hampered by screens that yield key myogenic regulators as candidates and a lack of mechanistic knowledge regarding their effects on *DUX4*. To uncover more specific targets, we performed a candidate-based screen which identified several epigenetic facilitators of *DUX4* expression in primary FSHD myocytes, including the chromatin remodeling factor BAZ1A. Here, we used a compound that we recently identified as a BAZ1A inhibitor and potent *DUX4* suppressor to interrogate the role of BAZ1A at the FSHD locus. Our data suggest a model in which BAZ1A binds to D4Z4 in FSHD muscle, changing the chromatin landscape of the array. BAZ1A binding leads to reduced occupancy of the HP1α repressor, increased occupancy of the p300 coactivator, and increased H3K27 acetylation, promoting transcription of both *DUX4* and the long non-coding RNA *DBE-T* from the disease locus. *DBE-T,* in turn, recruits the histone methyltransferase ASH1L, which establishes H3K36 methylation in *cis,* further promoting *DUX4* transcription. BAZ1A inhibition disrupts this powerful feed-forward loop, supporting the development of more metabolically stable inhibitors.

**GRAPHICAL ABSTRACT:** 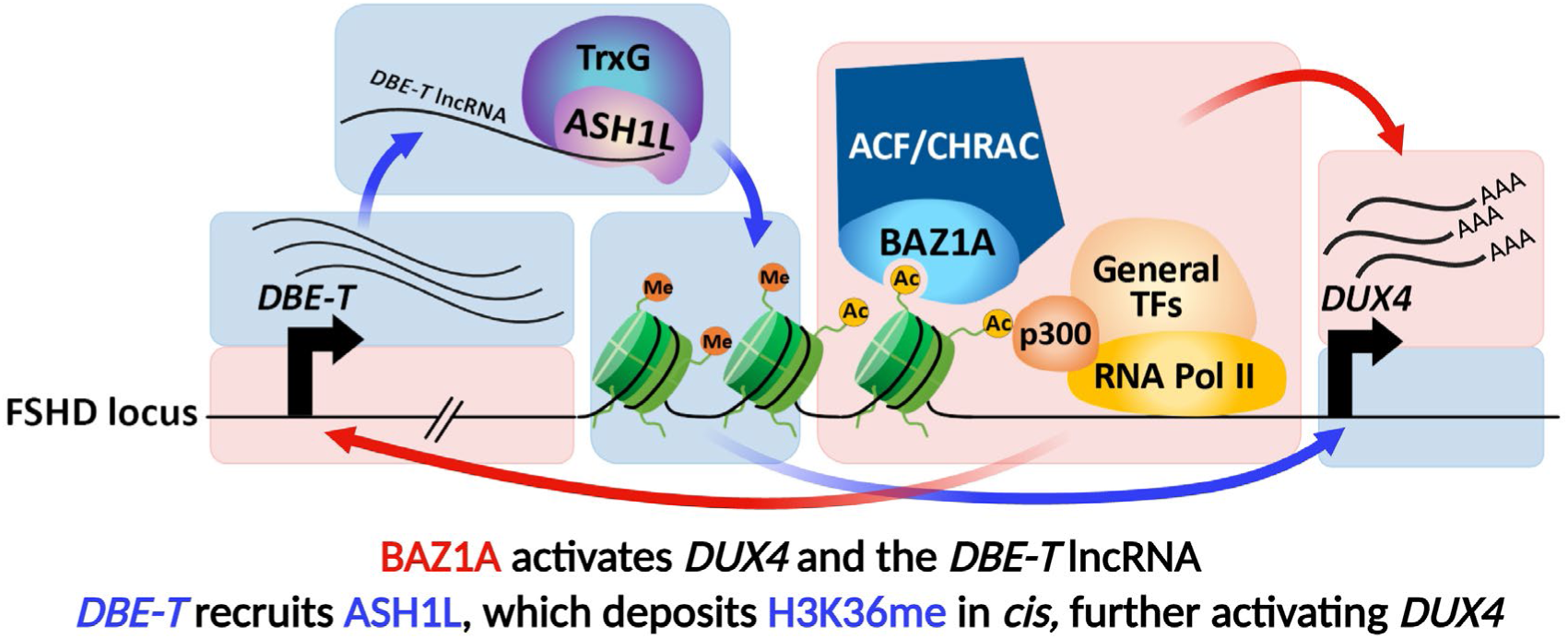

## INTRODUCTION

Facioscapulohumeral muscular dystrophy (FSHD, OMIM: 158900, 158901) is one of the most prevalent myopathies (1,2) and one that has generated considerable therapeutic interest over the past decade (3–7). All forms of the disease are caused by epigenetic dysregulation of the D4Z4 macrosatellite repeat array at chromosome 4q35 (Figure 1), which is normally silenced in adult somatic cells (8,9). FSHD1, the most common form of the disease, is linked to contractions at this array (10–12), resulting in chromatin relaxation. FSHD2 is caused by mutations in proteins that maintain epigenetic silencing of the D4Z4 array, leading to a similar chromatin relaxation (13–15). In both forms of the disease, loss of epigenetic repression leads to aberrant expression of the Double Homeobox 4 (*DUX4*) retrogene in skeletal muscle (16). DUX4, a cleavage-stage pioneer transcription factor, activates an early embryonic program, which causes pathology when misexpressed in adult skeletal muscle (9,17,18). Although an intact *DUX4* open reading frame resides in every D4Z4 repeat unit in the macrosatellite array (19), only the full-length *DUX4* mRNA *(DUX4-fl)* encoded by the distal-most repeat is stably expressed and translated due to the polyadenylation signal (PAS) residing in an exon distal to the array in disease-permissive alleles (16,20).

**Figure 1.**
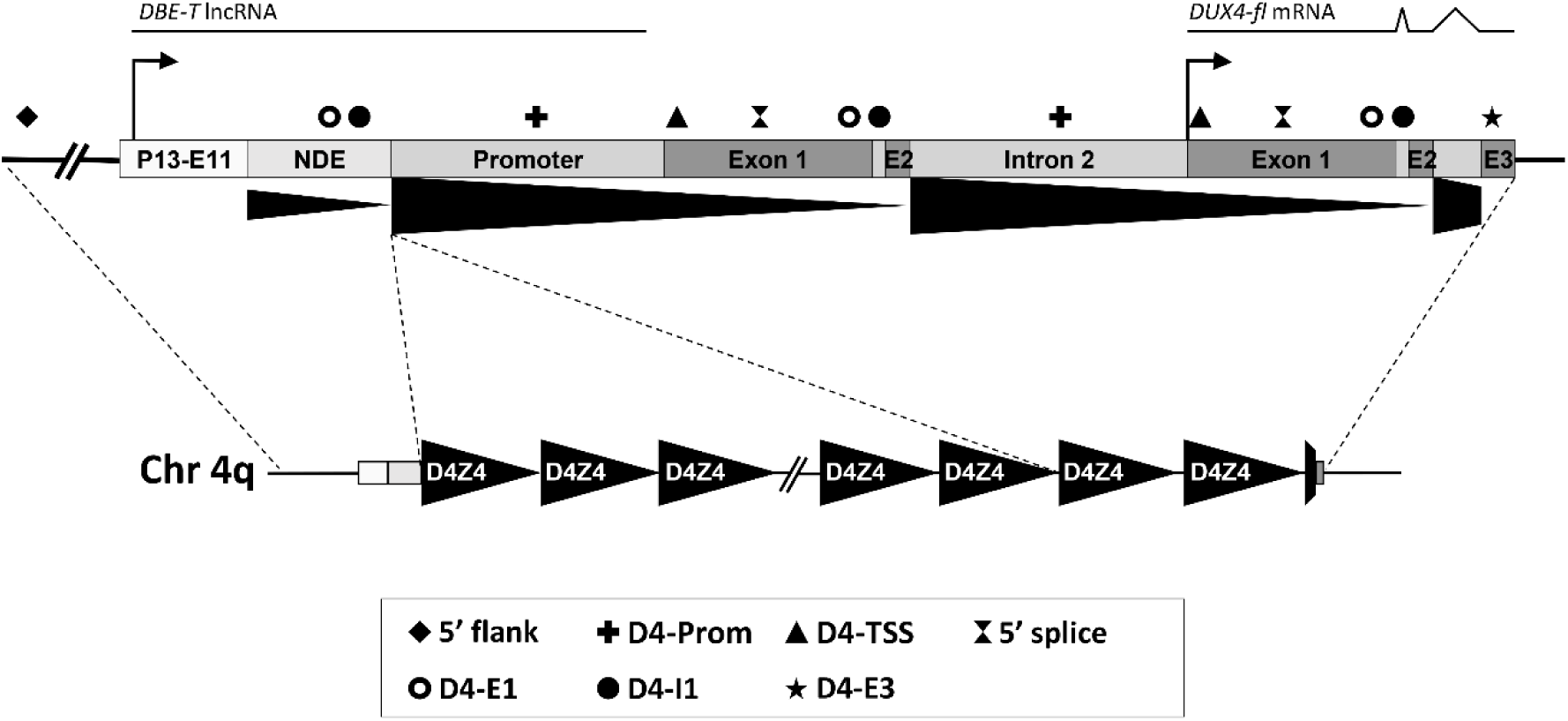
The FSHD disease locus. Schematic diagram of the FSHD locus at chromosome 4q35. The two distal-most repeat units are depicted above the D4Z4 macrosatellite array, with positions of the *DBE-T* lncRNA and pathogenic *DUX4-fl* mRNA indicated. The p13-E11 diagnostic probe region (21,22) and the non-deleted element (NDE) (8,23) lie proximal to the array. *DUX4* exons 1 and 2 are located within each repeat unit, and exon 3 lies in the distal subtelomeric sequence. In FSHD skeletal myocytes, *DUX4-fl* mRNA from the distal repeat is stabilized by a polyadenylation signal in exon 3 that is present in disease-permissive haplotypes. Regions amplified in ChIP assays (symbols) are: *DUX4* upstream flanking region (5’ flank), promoter (D4-Prom), translation start site (D4-TSS), 5’ splice junction (5’ splice), exon 1 (D4-E1), intron 1 (D4-I1), and exon 3 (D4-E3). Part of *DUX4* sequence is duplicated in the NDE.

While the repressive mechanisms that silence D4Z4 macrosatellite arrays in the healthy state have been well-characterized (9,24–26), much less is known about the factors and pathways that promote aberrant activation of *DUX4-fl* from a permissive array in FSHD muscle. These are of particular interest in the search for more specific targets for drug development. While small molecules remain the gold standard for therapeutics, only the p38 inhibitor losmapimod has been in clinical trials for FSHD (NCT04003974; NCT05397470). This repurposed cardiovascular drug failed its primary endpoint in Phase 3 and has always raised concerns regarding the detrimental effects of chronically targeting a key regulator of muscle biology in dystrophic muscles (27). Additionally, the mechanism(s) by which p38 facilitates *DUX4-fl* expression - and thus, the mechanism(s) by which losmapimod represses *DUX4-fl* - remain unknown. It is clear that more specific and better characterized targets for small molecule development in FSHD are needed.

Using a candidate gene knockdown approach, we identified several key activators of *DUX4* as potential therapeutic targets (28), including ASH1L (Absent, Small, or Homeotic 1 Like), BAZ1A (bromodomain adjacent to zinc finger domain 1A; ACF1), and SMARCA5 (SWI/SNF related matrix associated actin dependent regulator of chromatin subfamily A member 5; SNF2H) (28). ASH1L, a member of the Trithorax group of epigenetic activators, is a histone methyltransferase previously identified as an activator of *DUX4* (29). ASH1L is recruited to the FSHD locus by the *cis*-acting long non-coding RNA (lncRNA) *DBE-T* (D4Z4 Binding Element-Transcript) to establish active histone modifications (29). By contrast, the mechanism(s) by which SMARCA5 and BAZ1A activate *DUX4* in FSHD myocytes are not known. Both factors are components of the ACF and CHRAC chromatin remodeling complexes (30,31). Although non-enzymatic, BAZ1A plays a key regulatory role, controlling the directionality, efficiency, and specificity of the SMARCA5 ATPase, which utilizes ATP hydrolysis to remodel nucleosomes, regulating access to chromatin (32,33). As opposed to SMARCA5, which is a critical regulator of numerous cellular processes, BAZ1A has a much more limited role (34) and represents a relatively specific target for therapeutic development.

Here, we used a compound that we recently identified as a BAZ1A inhibitor and potent suppressor of *DUX4* (35) to interrogate the role of BAZ1A in regulating chromatin dynamics and transcription at the FSHD locus.

## MATERIALS AND METHODS

### Cell lines and cell culture

Primary human myogenic cells derived from biceps muscles of FSHD1 patients were obtained from the Sen. Paul D. Wellstone FSHD cell repository housed at the University of Massachusetts Medical School, as described (36). Primary myoblasts from unrelated FSHD1 patients (09Abic, 17Abic) or unaffected first-degree relatives (09Ubic, 17Ubic) were propagated in HMP growth medium [Ham’s F-10 medium (Corning^TM^ Cellgro^TM^, #10-070-CV) supplemented with 20% characterized FBS (Hyclone), 1% chick embryo extract (USBiological, #C3999), 1.2 mM CaCl_2_, and 1% antibiotics/antimycotics (Corning^TM^ Cellgro^TM^, #30-004-CI)] on plates coated with 0.1% gelatin (Sigma-Aldrich, #G1890). The immortalized 15Abic human skeletal muscle cell line (15Ai) was originally derived from the biceps muscle of an individual with FSHD (37,38). 15Ai myoblasts were expanded and cultured in RoosterNourish™-MSC medium (RoosterBio) on 0.1% gelatin-coated plates.

For differentiation, immortalized myoblasts were grown to confluence in 0.1% gelatin-coated plates and switched to NbActiv4 differentiation medium (Brainbits /TransnetYX Tissue, #Nb4-500) after a phosphate-buffered saline (PBS, Corning^TM^ Cellgro^TM^, #20-031-CV) wash. Dimethyl sulfoxide (DMSO, Sigma-Aldrich, #D1435) or compound C06 (previously dissolved in DMSO at 10 mM stock concentration) was added at the desired concentration and maintained for 4 days with daily media changes until harvesting. Upon reaching confluence, primary myoblasts were differentiated in Dulbecco’s Modified Eagle Medium (DMEM, Corning^TM^ Cellgro^TM^, #10-013-CV) containing 2% horse serum (Sigma-Aldrich, #H1270) and fed with fresh differentiation media every other day until harvesting or analysis.

### Lentiviral (LV) infections

LV particles expressing shRNAs against human *BAZ1A* (TRCN0000034279 and TRCN0000034280), human *ASH1L* (TRCN0000016169), control shRNA (TRCNS control), or an empty vector control (SHC001V) were generated from expression plasmids (MISSION shRNA library, MilliporeSigma) as described (28). For viral transduction, all myoblast lines were grown to confluence, then immortalized myoblasts were switched to differentiation conditions while primary myoblasts were allowed to self-differentiate in growth medium. All cells were subjected to two rounds of infection with LV supernatants and harvested 4 days later, as described (28).

### RNA extraction, reverse transcription, and quantitative real-time polymerase chain reaction (RT-qPCR)

Total RNAs were isolated from myoblasts or differentiated myotubes using TRIzol reagent (Ambion Life Technologies, #15596018) and purified using RNeasy MinElute Cleanup Kit (QIAGEN, #74204) according to the manufacturer’s instructions. Total RNAs (2 µg) were used for cDNA synthesis using Superscript III Reverse Transcriptase (Invitrogen, #56575), and 200 ng cDNA was used for SYBR green based qPCR analysis using the BioRad CFX96^TM^ Real-Time System (C1000 Touch^TM^ Thermal Cycler) as described (39). Oligonucleotide primer sequences are provided in Supplementary Table S1. *DBE-T* expression was analyzed in 09Abic and 09Ubic muscle cells, as described (29).

### Chromatin immunoprecipitation (ChIP)

ChIP assays were performed with C06-treated or shRNA-treated differentiated myotubes using the Fast ChIP method (40) as described (41). Cells were cross-linked by 1% formaldehyde (Sigma-Aldrich, #F1635) in DMEM for 10 minutes at room temperature followed by addition of 125 mM glycine in DMEM to stop the cross-linking. Cells were washed with ice-cold PBS supplemented with protease inhibitors, harvested with IP buffer [1% Triton X-100, 0.5% NP-40, 5 mM EDTA, 50 mM Tris-pH7.5 and 150 mM NaCl], then dounce-homogenized on ice for 10 strokes. Nuclei were pelleted by centrifugation at 12000x*g,* then resuspended in ice-cold IP buffer. Chromatin was sonicated in an ice bath using a Branson Digital Sonifier (EDP No. 100-132-885R, Branson Ultrasonics Corporation, CA, USA) at 65% amplitude (12 cycles of 15-sec pulses with 1 min and 45-sec rest after each pulse). 4% of the lysates were saved as input control. Chromatin was immunoprecipitated using 2 µg of specific antibodies against BAZ1A (Bethyl Laboratories, #A301-318A), p300 (Abcam, #ab14984), H3K27Ac (Abcam, #ab4729), HP1α (Abcam, #ab77256), SMARCA5 (Bethyl Laboratories, #A301-017A), ASH1L (Bethyl Laboratories, #A301-749A), H3K36me3 (Abcam, #ab9050), or H3K36me2 (Abcam, #ab9049). The immuno-complexes were collected using Protein A agarose beads (Cytiva, nProtein A Sepharose^TM^ 4 Fast Flow, #17528001), and washed 5X with IP buffer. Crosslinking was reversed using a 10% Chelex 100 Resin (BioRad, #142-1253) at 95°C for 10 minutes followed by digesting with 0.2 mg/ml of Proteinase K (Invitrogen, #100005393) at 55°C in a thermal mixer (1000 rpm) for 4 hours. Proteinase K was inactivated at 95°C for 10 minutes. Quantitative real-time PCR was performed as described (41). Oligonucleotide primer sequences are provided in Supplementary Table S1.

### Statistical analysis

All experiments were performed using at least three biological replicates. Data was expressed as mean + standard error of the mean (SEM) and was analyzed by unpaired, two-tailed Student’s *t*-test using GraphPad Prism 9.0 (GraphPad Software, Inc. La Jolla, CA, USA).

## RESULTS

We first determined the endogenous expression levels of *BAZ1A* in FSHD and healthy myocytes, and correlated these with expression of *DUX4-fl*. While *DUX4-fl* was only detectable in differentiated FSHD myocytes, *BAZ1A* levels were equivalent in the FSHD vs. healthy state, and in the differentiated vs. proliferating state (Figure 2A). We then verified binding of BAZ1A at the FSHD locus using chromatin immunoprecipitation (ChIP) assays. Interestingly, BAZ1A was enriched at the *DUX4* locus in FSHD, but not healthy, myotubes (Figure 2B).

**Figure 2.**
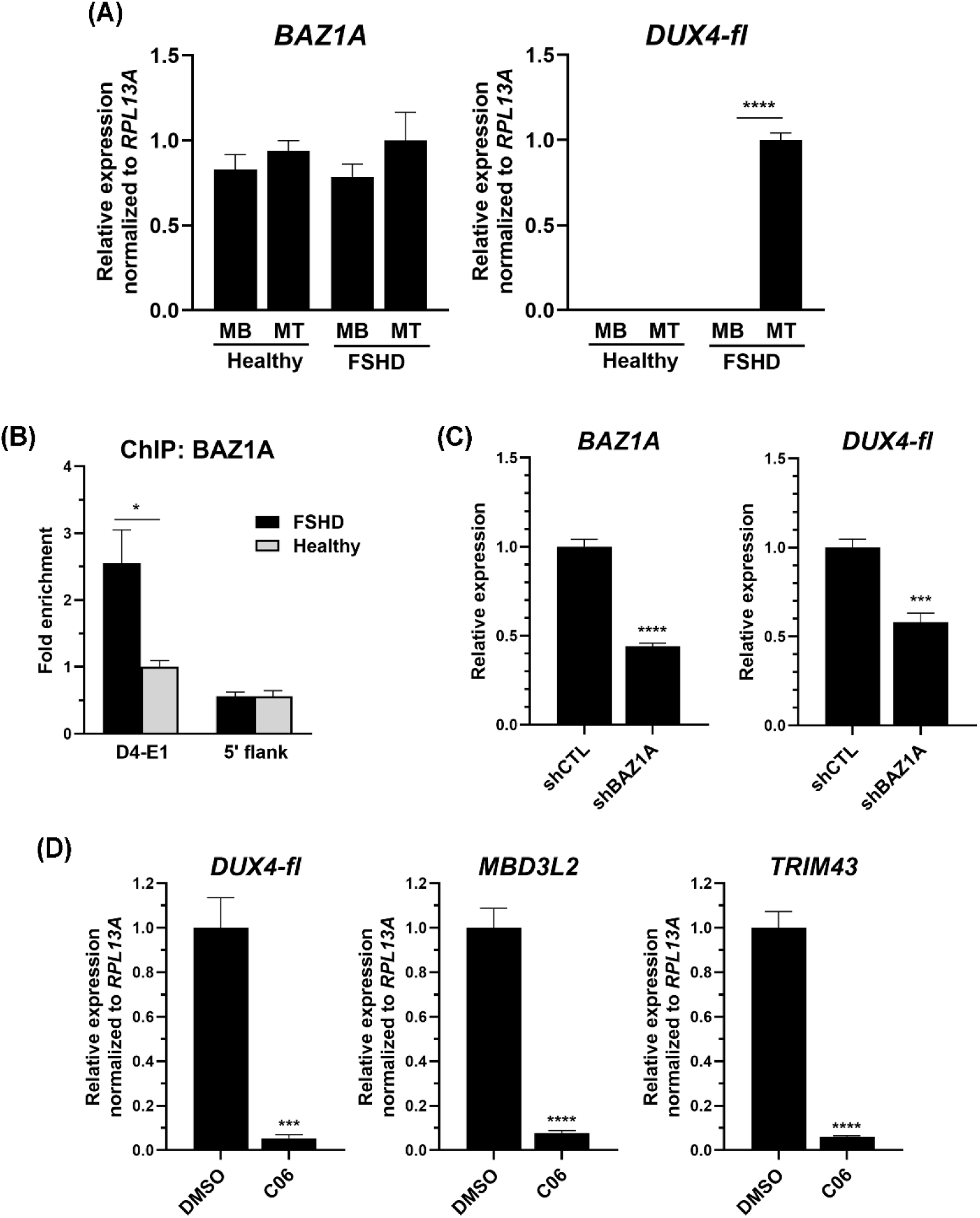
BAZ1A binds D4Z4 chromatin in FSHD myotubes, facilitating *DUX4-fl* expression. **(A)** Primary muscle cells from an FSHD patient or a healthy relative were grown and harvested as proliferating myoblasts (MB) or differentiated myotubes (MT). Expression levels of *BAZ1A* and *DUX4-fl* were assessed by RT-qPCR and normalized to levels of *RPL13A.* **(B)** ChIP assays were performed on differentiated FSHD and healthy myotubes using antibodies specific for BAZ1A, and analyzed by qPCR using primers to *DUX4* exon 1 (D4-E1) or a control region upstream of *DUX4* (5’ flank). Data are presented as the relative fold enrichment normalized to enrichment at D4-E1 in healthy cells. **(C)** FSHD myotubes were infected with lentiviral vectors encoding shRNAs against *BAZ1A* (shBAZ1A) or a control sequence (shCTL). Expression levels of *BAZ1A* and *DUX4-fl* were assessed by RT-qPCR and normalized to *RPL13A.* **(D)** FSHD myotubes were treated with 50 nM C06 or DMSO control. Expression levels of *DUX4-fl* and DUX4-FL target genes *MBD3L2* and *TRIM43* were assessed by RT-qPCR and normalized to *RPL13A*. For all panels, data are plotted as the mean + standard error of the mean (SEM) of at least three independent experiments. \**p* < 0.05, \*\*\**p* < 0.001, and \*\*\*\**p* < 0.0001 by unpaired *t* test.

Inducing efficient gene knockdowns in skeletal myocytes using RNA interference systems, such as shRNAs, requires serial lentiviral (LV) infections as opposed to the more facile lipid-mediated transfections used in other cell types. In addition, *DUX4-fl* expression is closely linked to myogenic differentiation and cellular stress; thus, any factor that affects these pathways (such as the stress induced by viral transduction) can alter *DUX4-fl* levels artifactually, rather than specifically. With this in mind, we sought a more efficient and specific approach for manipulating BAZ1A activity.

In a recent study, we used an artificial intelligence (AI)-based approach to identify small molecules predicted to interact with the BAZ1A bromodomain, and validated these compounds for their ability to reduce *DUX4-fl* expression in FSHD myocytes (35). This study identified the C06 compound as a potent suppressor of *DUX4-fl* (35). Interestingly, while C06 exhibits binding to BAZ1A, it can also inhibit multiple kinases, including p38α, a known activator of *DUX4-fl* and drug target in FSHD (35). However, RNA-seq assessment of FSHD myocytes treated with C06 showed that while DUX4-FL targets were returned to a healthier expression profile, expression of myogenic factors was not significantly altered and there were no major effects on global gene expression (35). This suggests that although C06 is capable of inhibiting multiple pathways, its effects, especially at low concentrations, are fairly specific. To validate a more recent batch of C06 with the cells used in the current study, we used RT-qPCR to compare the amount of *DUX4-fl* repression achieved in FSHD myocytes following treatment with C06 vs. with shRNAs specific to *BAZ1A.* While shRNA knockdown of *BAZ1A* yielded a modest ∼45% repression of *DUX4-fl* (Figure 2C), treatment with C06 resulted in a striking 95% repression of *DUX4-fl* and DUX4-FL targets *MBD3L2* and *TRIM43* (Figure 2D).

To gain insight into the role of BAZ1A at the *DUX4* locus, we performed ChIP assays using chromatin from FSHD myocytes treated with C06. BAZ1A inhibition with C06 reduced occupancy of BAZ1A and the transcriptional activator p300 across the *DUX4* gene body (Figure 3A-B). p300 is a histone acetyltransferase that catalyzes acetylation of histone H3 lysine 27 (H3K27Ac), a marker of transcriptionally active chromatin, which was also reduced at *DUX4* in response to C06 treatment (Figure 3C). Additionally, occupancy of heterochromatin protein 1 alpha (HP1α) was increased at *DUX4* (Figure 3D). While HP1γ is the predominant isoform of this repressor found at D4Z4 chromatin in FSHD myocytes (42), we have found that HP1α is often enriched at the *DUX4* locus following perturbations that return the region to a healthier state (43).

**Figure 3.**
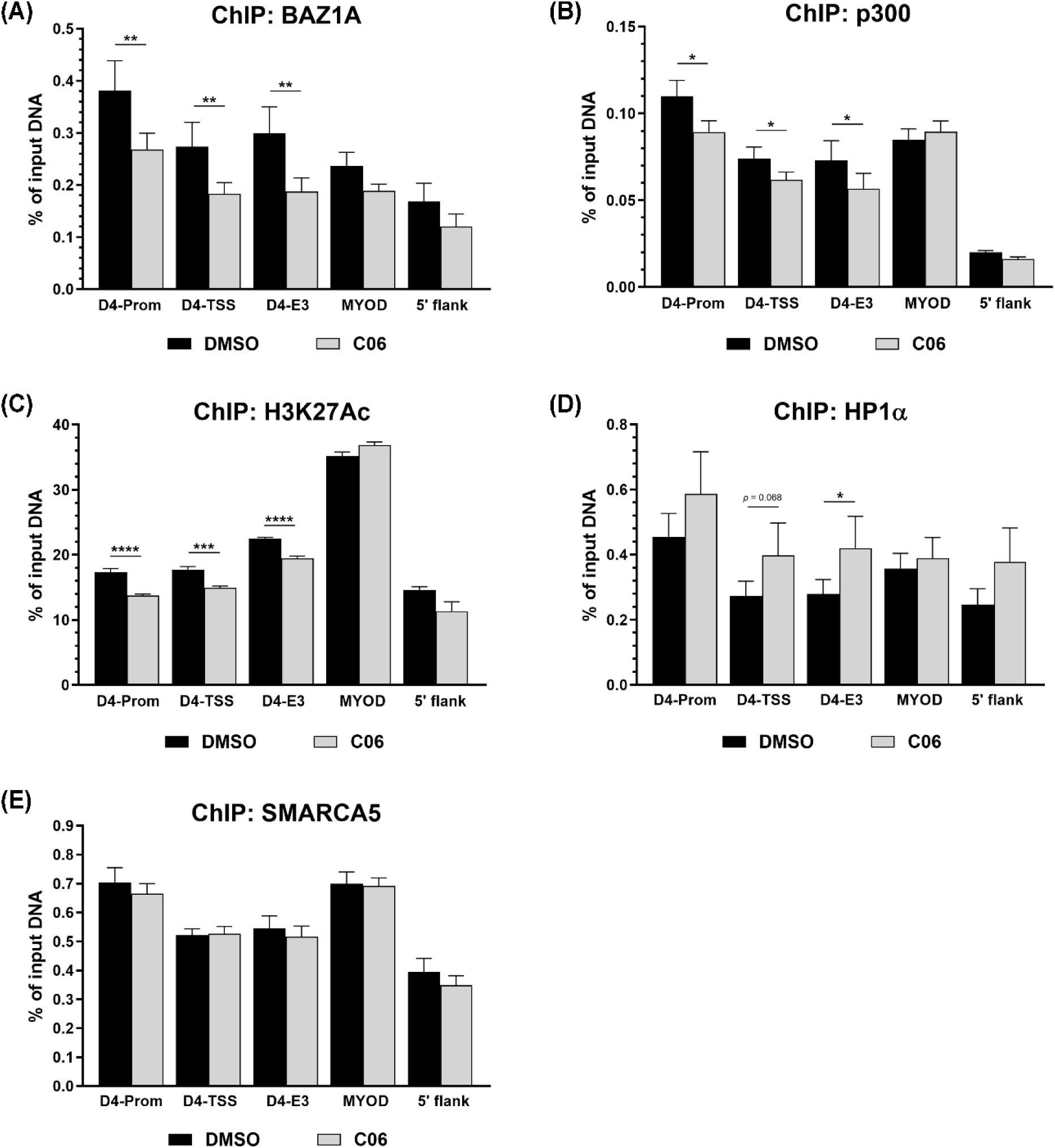
BAZ1A promotes active chromatin at D4Z4 in FSHD myotubes. **(A-E)** ChIP assays were performed on differentiated FSHD myotubes treated with 50 nM C06 or DMSO control using antibodies specific for BAZ1A, p300, acetylated histone H3 lysine 27 (H3K27Ac), heterochromatin protein 1α (HP1α), or SMARCA5, and analyzed by qPCR using primers to the *DUX4* locus (Figure 1). *MYOD1* was used for comparison with a highly transcriptionally active gene. Data was normalized to total input. Each bar represents the average of at least three independent ChIP experiments (mean + SEM). \**p* < 0.05, \*\**p* < 0.01, \*\*\**p* < 0.001, and \*\*\*\**p* < 0.0001 by unpaired *t* test.

In the context of chromatin remodeling, SMARCA5 and BAZ1A bind independently of each other at DNA lesions (44), but less is known about their binding mechanisms at sites of gene regulation. At the *DUX4* locus, we found that BAZ1A inhibition had no effect on SMARCA5 occupancy (Figure 3E). By contrast, shRNA-mediated knockdown of *SMARCA5* reduced both SMARCA5 and BAZ1A occupancy at *DUX4* (Figure 4A-B). This suggests that SMARCA5 can bind D4Z4 chromatin independent of BAZ1A, but BAZ1A binding may be dependent on SMARCA5. As with BAZ1A inhibition, *SMARCA5* knockdown reduced p300 occupancy and H3K27Ac enrichment at *DUX4* (Figure 4C-D), although occupancy of HP1α was unaffected (Figure 4E). Importantly, *SMARCA5* knockdown had no significant effect on *BAZ1A* expression (Figure 4F), indicating that the lower levels of BAZ1A at *DUX4* are due to reduced occupancy rather than decreased expression. Interestingly, either C06 treatment or *SMARCA5* knockdown had no effect on occupancy of any of the tested factors at the *MYOD1* locus, except that levels of H3K27Ac were increased, rather than decreased, in response to *SMARCA5* knockdown (Figures 3 and 4). This is in stark contrast to effects on D4Z4 chromatin, and it suggests that the dynamics of BAZ1A and SMARCA5 at the transcriptionally active *MYOD1* gene are very different from those at the aberrantly active *DUX4* gene.

**Figure 4.**
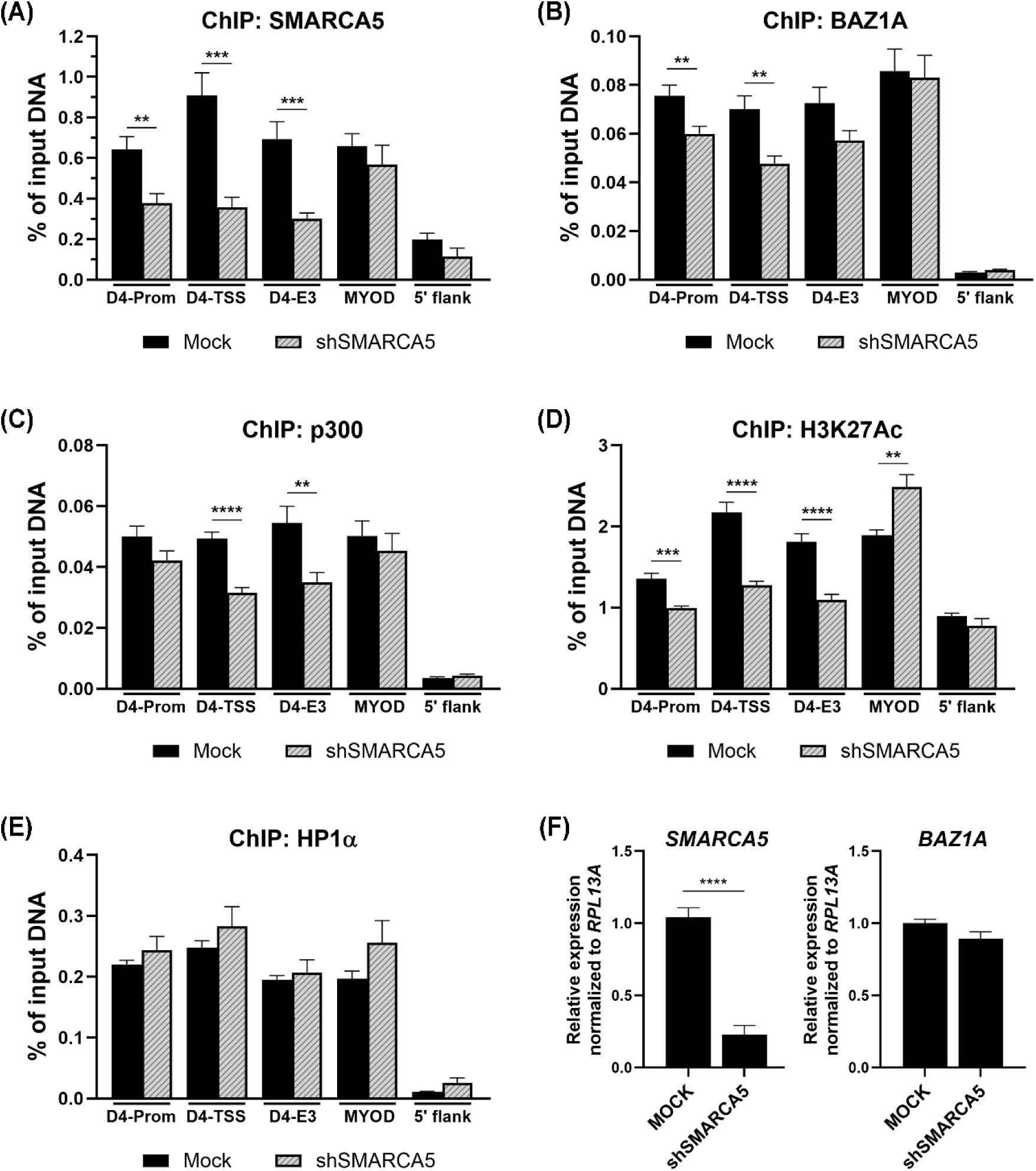
SMARCA5 facilitates binding of BAZ1A and establishment of active chromatin at D4Z4 in FSHD myotubes. **(A-E)** Differentiated FSHD myotubes were transduced with lentiviral vectors encoding shRNAs against *SMARCA5* (shSMARCA5) or mock infected. ChIP assays were performed using antibodies specific for SMARCA5, BAZ1A, p300, acetylated histone H3 lysine 27 (H3K27Ac), or heterochromatin protein 1α (HP1α), and analyzed by qPCR using primers to the *DUX4* locus (Figure 1). *MYOD1* was used for comparison with a highly transcriptionally active gene. Data was normalized to total input. **(F)** FSHD myotubes were transduced as above. Expression levels of *SMARCA5* and *BAZ1A* were assessed by RT-qPCR and normalized to *RPL13A.* For all panels, each bar represents the average of at least three independent experiments (mean + SEM). \*\**p* < 0.01, \*\*\**p* < 0.001, and \*\*\*\**p* < 0.0001 by unpaired *t* test.

Chromatin remodeling complexes such as ACF play diverse, context-dependent roles in gene regulation. In addition to mediating Polycomb repression (45), the ACF complex and its individual components can establish hypersensitive sites and activate/repress gene expression in concert with certain transcription factors (46–51). Thus, we considered the possibility that BAZ1A might be exerting its positive effects on *DUX4* through another D4Z4 regulator. A likely candidate for this is ASH1L, the only factor that has been shown to directly activate the FSHD disease locus (29). A histone methyltransferase that deposits H3K4me3 and H3K36me2/3 at its gene targets, ASH1L is the mammalian homologue of the Drosophila Trithorax group protein that counteracts Polycomb-mediated gene silencing (52–54). It is recruited proximal to the D4Z4 array by the *DBE-T* lncRNA, resulting in methylation at H3K36, de-repression of the disease locus, and aberrant *DUX4* expression in FSHD myocytes (29). ASH1L also emerged as a *DUX4* activator from our candidate-based screen that yielded BAZ1A and SMARCA5 as potential therapeutic targets (28).

We confirmed that occupancy of ASH1L is enriched at *DUX4* in FSHD vs. healthy myocytes (Figure 5A), and that *ASH1L* knockdown reduces the H3K36me3 mark across the locus (Figure 5B). To determine whether BAZ1A is required for ASH1L recruitment, we treated cells with C06, and found a striking decrease in levels of *DBE-T* (Figure 5C). Consistent with this, BAZ1A inhibition decreased occupancy of ASH1L and levels of H3K36me2/3 across the *DUX4* locus (Figure 5D-F). Taken together, these results demonstrate that BAZ1A is required for expression of *DBE-T,* which recruits ASH1L to the FSHD locus, promoting H3K36me deposition and transcription of *DUX4-fl*.

**Figure 5.**
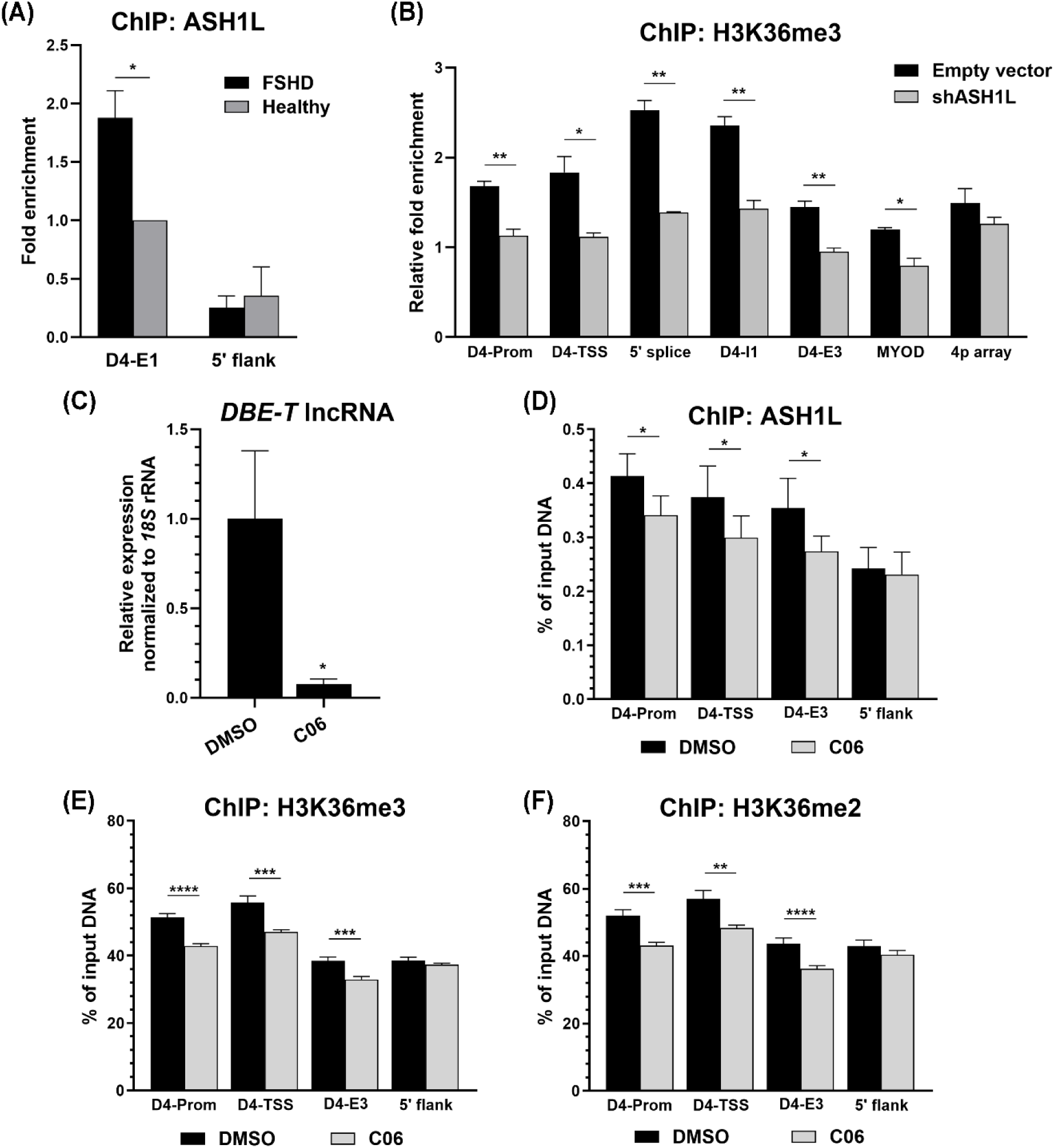
BAZ1A facilitates expression of the *DBE-T* lncRNA and recruitment of ASH1L to D4Z4 in FSHD myotubes. **(A)** ChIP assays were performed on differentiated FSHD and healthy myotubes using antibodies specific for ASH1L, and analyzed by qPCR using primers to the *DUX4* locus (Figure 1). **(B)** Differentiated FSHD myotubes were transduced with lentiviral vectors encoding shRNAs against *ASH1L* (shASH1L) or empty vector. ChIP assays were performed using antibodies specific for tri-methylated histone H3 lysine 36 (H3K36me3), and analyzed by qPCR using primers to the *DUX4* locus. *MYOD1* was used for comparison with a highly transcriptionally active gene, and the chromosome 4p macrosatellite array (4p array) was used for comparison with a transcriptionally silent region. **(C)** Expression of the *DBE-T* lncRNA was assayed by RT-qPCR in differentiated FSHD myotubes treated with 50 nM C06 or vehicle (DMSO) and normalized to *RPL13A*. **(D-F)** FSHD myotubes were treated with 50 nM C06 or DMSO. ChIP assays were performed using antibodies specific for ASH1L (D), H3K36me3 (E), or H3K36me2 (F), and analyzed by qPCR using primers to the *DUX4* locus. For all ChIP experiments, data was normalized to total input. For all panels, each bar represents the average of at least three independent experiments (mean + SEM). \**p* < 0.05, \*\**p* < 0.01, \*\*\**p* < 0.001, and \*\*\*\**p* < 0.0001 by unpaired *t* test.

## DISCUSSION

While much is known regarding normal mechanisms of D4Z4 repression, mechanisms of *DUX4* activation in the FSHD disease state are underexplored, especially considering that these represent potential targets for traditional drug development. BAZ1A and SMARCA5 emerged as two key activators of *DUX4-fl* in our previous screen (28). While both factors are part of the ACF and CHRAC chromatin remodeling complexes (34), SMARCA5 is also a critical component of at least six other complexes involved in transcriptional regulation, ribosomal gene expression, DNA repair, DNA replication, and the DNA damage response (34). Thus, BAZ1A represents a more specific target than SMARCA5 for therapeutic development, and inhibition of BAZ1A offers a way to examine the interplay of these two factors at the FSHD disease locus with minimal disruption of other complexes and pathways.

The BAZ1A inhibitor C06, identified in our previous study as a potent repressor of *DUX4-fl* (35), provides a useful new tool for interrogating BAZ1A function. Previously, affecting the expression or activity of BAZ1A required RNAi or CRISPR-based approaches, which necessitate viral transduction in skeletal myocytes. This results in the induction of many viral-response genes, complicating data interpretation (43,55). In our previous study, the RNA-seq gene ontology analysis confirmed that shRNA knockdown of *BAZ1A* induced numerous changes in the muscle transcriptome, including the induction of many viral-response genes from the transduction (35). By contrast, treatment with C06 at a range of concentrations yielded only a small number of DEGs, many of which are DUX4 targets (35).

Thus, we used C06 in the current study to investigate the mechanisms by which BAZ1A contributes to *DUX4-fl* expression in FSHD skeletal myocytes. Taken together, our data suggests a model in which BAZ1A binds to D4Z4 and changes the chromatin landscape of the array (Figure 6). BAZ1A binding is associated with reduced occupancy of the HP1α repressor and increased H3K27 acetylation and occupancy of the p300 coactivator, promoting transcription of both *DUX4-fl* and the cis-acting lncRNA *DBE-T*. *DBE-T,* in turn, recruits ASH1L, which establishes H3K36 methylation, further promoting *DUX4-fl* transcription (29).

**Figure 6.**
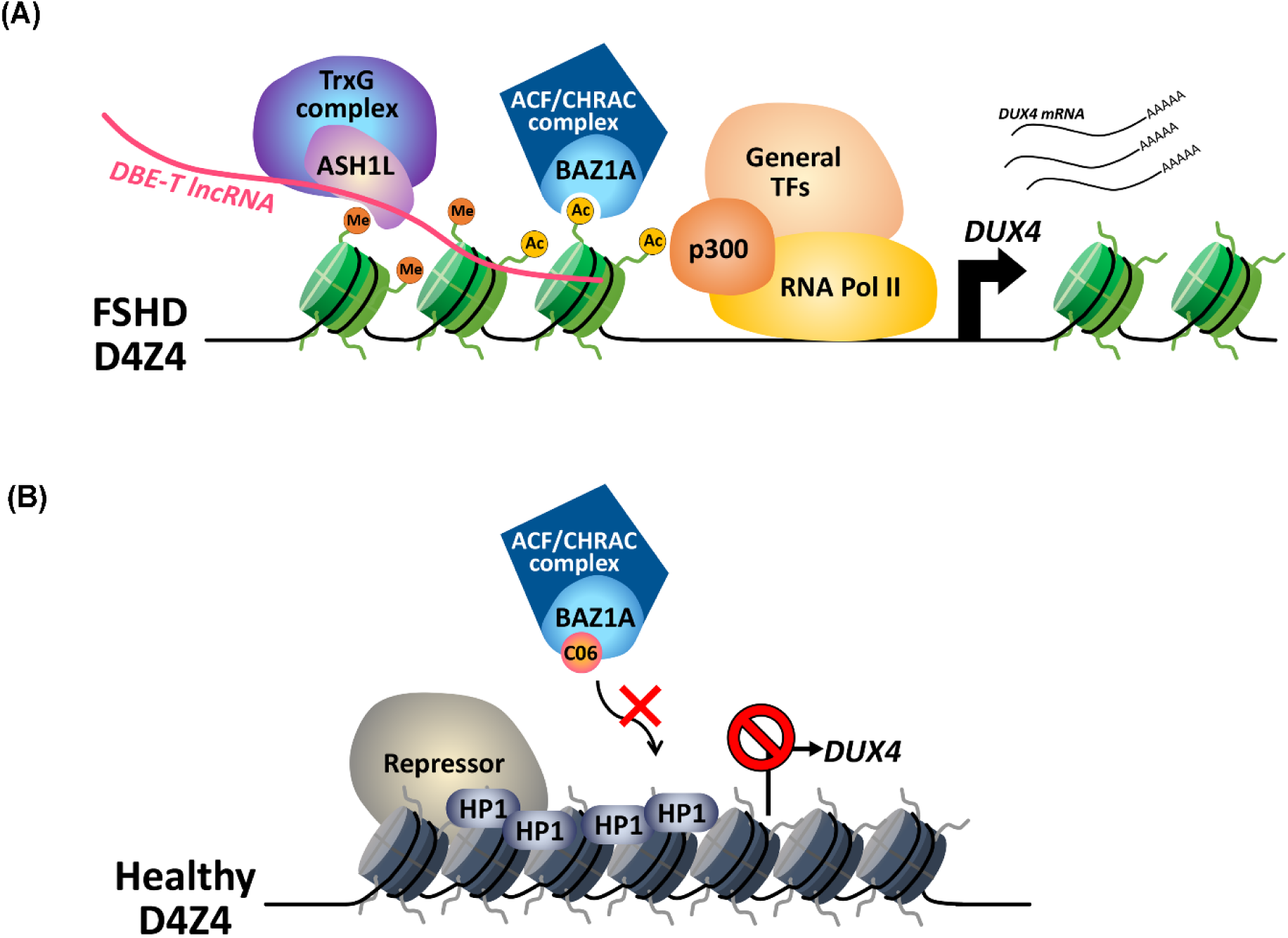
Model of BAZ1A regulation of the FSHD locus. **(A)** Our data supports a model in which BAZ1A binds to the D4Z4 array in FSHD myotubes, changing the chromatin landscape of the array. BAZ1A binding is associated with reduced occupancy of the HP1α repressor, increased occupancy of the p300 coactivator, and increased H3K27 acetylation, promoting transcription of both *DUX4-fl* and the lncRNA *DBE-T* from the locus. *DBE-T,* in turn, recruits the histone methyltransferase ASH1L, which establishes H3K36 methylation in *cis,* further promoting *DUX4-fl* transcription. **(B)** Inhibition of BAZ1A interferes with its binding to D4Z4, leading to a more repressive chromatin state and reduced transcription of *DBE-T.* This results in decreased recruitment of ASH1L to the locus, and consequently reduced transcription of *DUX4-fl.* ACF, ATP-utilizing chromatin assembly and remodeling factor; ASH1L, ASH1-like histone lysine methyltransferase; BAZ1A, Bromodomain Adjacent To Zinc Finger Domain 1A; CHRAC, chromatin accessibility complex; *DBE-T* lncRNA, *D4Z4 Binding Element Transcript* long non-coding RNA; HP1, Heterochromatin Protein 1; RNA Pol II, RNA polymerase II; TFs, transcription factors; TrxG, Trithorax-group.

Many small-molecule bromodomain inhibitors are currently in development as epigenetic targeted therapies for cancers and inflammatory disorders (56–61). A number of these target the bromodomain and extra-terminal (BET) family, whose members share a common domain architecture featuring two amino-terminal bromodomains (59). While BET inhibitors are potent repressors of *DUX4* expression in FSHD myocytes (62), their targets have not been demonstrated to function directly at the FSHD locus. Instead, BET family members occupy many transcriptionally active promoters, including those of critical myogenic regulators (63), raising specificity concerns regarding the use of BET inhibitors as FSHD therapeutics. Similarly, p38α, the target of losmapimod, which recently failed in a Phase 3 clinical trial for FSHD (NCT05397470) despite promising preclinical data (64), has no known direct connection to the FSHD locus, and the mechanism by which it regulates *DUX4* is unknown. Additionally, it does not appear to regulate *DUX4* at the late stage of muscle differentiation (65). In contrast to this, we show in the current study that BAZ1A is directly involved in mediating the active state of D4Z4 chromatin, and that its activation of the *DBE-T* lncRNA provides positive feedback for *DUX4-fl* expression. As opposed to BET or p38 inhibition, BAZ1A inhibition has no significant effects on myogenic genes (28) and it suppresses *DUX4-fl* through a clearly elucidated mechanism: impeding BAZ1A binding to the disease locus, which reduces active chromatin changes and the subsequent expression of both *DUX4-fl* and the *DBE-T* lncRNA that promotes *DUX4-fl* expression.

Interestingly, as with many lncRNAs that recruit chromatin modifying complexes to genomic loci (66,67), *DBE-T* recruits both ASH1L and the epigenetic scaffold WDR5 to the *DUX4* locus in FSHD myocytes (68). While inhibition of WDR5 was also demonstrated to suppress *DUX4-fl* (68), this protein is a core component of HMT complexes, it is essential for development, and it is a critical regulator of numerous cellular processes (69). Even relatively specific WIN site antagonists of the interaction between WDR5 and the MLL/SET1 methyltransferase complexes cause a downregulation of protein synthesis genes (68,70). BAZ1A represents a more selective and potentially more powerful target for therapy, considering its more limited cellular role and the fact that it functions upstream of *DBE-T,* a locus-specific regulator.

Importantly, even the modest reduction of *BAZ1A* achieved by shRNA knockdown or CRISPR inhibition significantly reduced the expression of *DUX4-fl* (28). While therapeutic levels of DUX4 inhibition are unknown, data from clinically affected, asymptomatic, and non-manifesting FSHD subjects support the idea that any reduction in *DUX4* expression will have therapeutic benefit (36,39,71). Thus, even a partial reduction of BAZ1A expression or activity will likely be clinically beneficial. Efforts to engineer more metabolically stable compounds based on C06 are currently underway.

## SUPPLEMENTARY DATA

Supplementary Data is available at NAR Molecular Medicine online.

## AUTHOR CONTRIBUTIONS

NC, TIJ, PLJ, and CLH designed the study and analyzed data. NC and CLH performed experiments and wrote the manuscript. All authors edited and approved the final version of the manuscript.

## DATA AVAILABILITY

The authors declare that the data supporting the findings of this study are available within the main manuscript and its online supplementary materials.

