## Supplementary Figure S1 and Supplementary Table S1 for "BAZ1A promotes expression of *DUX4-fl* and its lncRNA activator *DBE-T* in facioscapulohumeral muscular dystrophy"

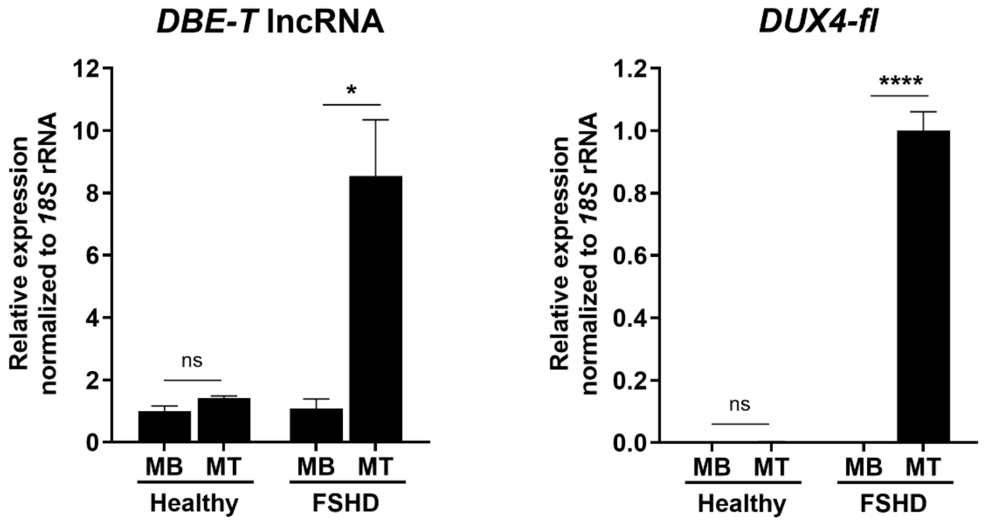


**Supplementary Figure S1.** ***DBE-T* lncRNA is exclusively expressed in differentiated FSHD myotubes.**

Relative expression levels of **(A)** the *DBE-T* lncRNA and **(B)** *DUX4-fl* mRNA were assessed by RT-qPCR in proliferating myoblasts (MB) and 6-day differentiated myotubes (MT) from an FSHD1 patient and unaffected family member. Data was normalized to levels of *18S* rRNA and plotted as the mean + standard error of the mean (SEM) of three independent experiments. $^*p < 0.05$ and $^{****}p < 0.0001$ by unpaired $t$ test.

**Supplementary Table S1. Oligonucleotide primers**

**qRT-PCR:**

| Primer name | Sequence | Reference |
|---|---|---|
| *BAZ1A*-F | 5′-CACCGAAAGCCGTTTGTGAG-3′ | (1) |
| *BAZ1A*-R | 5′-TCTACCCGTCACAGCACAAC-3′ | |
| *DUX4-fl*-F (prePCR) | 5′-GCTCTGCTGGAGGAGCTTTAGGA-3′ | |
| *DUX4-fl*-R (prePCR) | 5′-CGCACTGCTCGCAGGTCTGCWGGT-3′* | |
| *DUX4-fl*-nested-F | 5′-AGCTTTAGGACGCGGGGTTGGGAC-3′ | (1) |
| *DUX4-fl*-nested-R | 5′-GCAGGTCTGTWGGTACCTGG-3′* | |
| *MBD3L2*-F | 5′-GCGTTCACCTCTTTTCCAAG-3′ | (2) |
| *MBD3L2*-R | 5′-GCCATGTGGATTTCTCGTTT-3′ | |
| *TRIM43*-F | 5′-ACCCATCACTGGACTGGTGT-3′ | (2) |
| *TRIM43*-R | 5′-CACATCCTCAAAGAGCCTGA-3′ | |
| *RPL13A*-F | 5′-AACCTCCTCCTTTTCCAAGC-3′ | |
| *RPL13A*-R | 5′-GCAGTACCTGTTTAGCCACGA-3′ | |
| *SMARCA5*-F | 5′-TCAGGTCCGAGGATTAAACTG-3′ | |
| *SMARCA5*-R | 5′-ATATGAGGCCCAGGAATGTTT-3′ | |
| *DBE-T*-F | 5′-AGGCCTCGACGCCCTGGGTC-3′ | (3) |
| *DBE-T*-R | 5′-TCAGCCGGACTGTGCACTGCGGC-3′ | |
| *18S* rRNA-F | 5′-AGTAAGTGCGGGTCATAAGCT-3' | (1) |
| *18S* rRNA-R | 5′-CCTCACTAAACCATCCAATCGG-3' | |

*W=A or T

Underline: intentionally introduced mutation to ease primer dimer

**ChIP assay:**

| Primer name | Sequence | Reference |
|---|---|---|
| 5'flank-F | 5′-ATCTGGTTGTGGTAGTGTGC-3′ | (4) |
| 5'flank-R | 5′-TGAGGGTGTCTGAAAGAATG-3′ | |
| D4-Prom-F | 5′-CCTGTTGCTCACGTCTCTCC-3′ | (4) |
| D4-Prom-R | 5′-GTGGGGAGTCTGCAGTGTG-3′ | |
| D4-TSS-F | 5′-GACACCCTCGGACAGCAC-3′ | (4) |
| D4-TSS-R | 5′-GTACGGGTTCCGCTCAAAG-3′ | |
| 5'splice-F | 5′-CTGGTTTCAGAATCGAAGG-3′ | |
| 5'splice-R | 5′-CTGCCTGGCTCACGAAAG-3′ | |
| D4-E1-F | 5′-CGCAACCTCTCCTAGAAAC-3′ | |
| D4-E1-R | 5′-CAGAGCCCGGTATTCTTC-3′ | |
| D4-I1-F | 5′-CTCAGCGAGGAAGAATACCG-3′ | (2) |
| D4-I1-R | 5′-AGTCTCTCACCGGGCCTAGA-3′ | |
| D4-E3-F | 5′-CTGACGTGCAAGGGAGCT-3′ | (2) |
| D4-E3-R | 5′-CAGGTTTGCCTAGACAGCG-3′ | |
| *MYOD*-F | 5′-CGCCAGGATATGGAGCTACT-3′ | (5) |
| *MYOD*-R | 5′-CGGGTCGTCATAGAAGTCGT-3′ | |
| 4p array-F | 5′-TGGGAAATACCTGCTACGTG-3′ | (1) |
| 4p array-R | 5′-GTGACGATGACACGTTTGAG-3′ | |
